# Eco-evolutionary buffering: rapid evolution facilitates regional species coexistence despite local priority effects

**DOI:** 10.1101/040659

**Authors:** Meike J. Wittmann, Tadashi Fukami

## Abstract

Inhibitory priority effects, in which early-arriving species exclude competing species from local communities, are thought to enhance regional species diversity via community divergence. Theory suggests, however, that these same priority effects make it difficult for species to coexist in the region unless individuals are continuously supplied from an external species pool, often an unrealistic assumption. Here we develop an eco-evolutionary hypothesis to solve this conundrum. We build a metacommunity model in which local priority effects occur between two species via interspecific interference. Within each species there are two genotypes: one is more resistant to interspecific interference than the other, but pays a fitness cost for its resistance. Because of this trade-off, species evolve to become less resistant as they become regionally more common. Rare species can then invade some local patches and consequently recover in regional frequency. This “eco-evolutionary buffering” enables the regional coexistence of species despite local priority effects, even in the absence of immigration from an external species pool. Our model predicts that eco-evolutionary buffering is particularly effective when local communities are small and connected by infrequent dispersal.

## Introduction

There is now ample evidence that the effects that species exert on one another in a local habitat patch often depend on the order and initial abundance in which species arrive (Sutherland 1974; Drake 1991; Chase 2003). Known as priority effects (Slatkin 1974), such historical contingency in local community assembly is increasingly recognized as a major factor influencing species diversity (Fukami 2015). Specifically, recent research has suggested that local priority effects can enhance beta diversity, i.e., the variation in species composition among local communities, by driving communities onto divergent successional trajectories (e.g., Chase 2010; Martin and Wilsey 2012; Fukami and Nakajima 2013; Vannette and Fukami 2017).

For local priority effects to occur, patches must receive immigrants belonging to multiple species. This requirement can be easily met under the assumption that there is an external species pool. That is, immigrants entering local patches are drawn from a regional pool whose species composition is static and is not influenced by local community dynamics, as assumed by the classical theory of island biogeography (MacArthur and Wilson 1967). However, at large spatial and temporal scales, the regional pool consists of immigrants originating from other local patches (Mittelbach and Schemske 2015). In other words, the regional pool is not external, but instead internal *(sensu* Fukami 2005, 2015), as depicted by the metacommunity concept (Leibold et al.2004). To explain species diversity at these large scales, it is therefore necessary to understand how a diverse species pool can be maintained as a collective result of local community dynamics. This task is challenging when species engage in inhibitory priority effects, in which species that are initially common hinder colonization by competing species, a form of positive frequency dependence (Shurin et al. 2004). In many cases, species are likely to arrive at a newly created or disturbed patch in proportion to their regional abundances within the metacommunity. This correspondence between regional frequency and arrival probability can eventually result in regional extinction of all but one species (Taneyhill 2000; Shurin et al. 2004).

Thus, to maintain both local priority effects and a diverse regional pool of species, there has to be a mechanism that buffers species from regional extinction. Shurin et al. (2004) suggested that spatial environmental heterogeneity could be one such mechanism. In their model, patches differ in the relative rates of the supply of two essential resources. Two species could then engage in priority effects in patches with relatively balanced resource supply rates, whereas they exclude each other independently of initial composition in patches having more extreme supply rates. The extreme patches serve as refuges from which species can continue to disperse into patches where priority effects occur. In this sense, spatial refuges play a role qualitatively identical to that of an external species pool.

In this paper, we build a simple metacommunity model to suggest a new mechanism for the regional coexistence of species engaged in local inhibitory priority effects. The mechanism, which we call “eco-evolutionary buffering”, involves rapid evolution (*sensu* Hairston et al. 2005) of traits that determine how species interact. Previous studies of priority effects often assumed fixed species traits, but growing evidence suggests that traits often evolve at rates comparable to that of ecological population dynamics (Thompson 1998; Schoener 2011), which can then affect priority effects (Urban and De Meester 2009; Knope et al. 2012). For example, Urban and De Meester (2009) predicted that, given spatial environmental heterogeneity, rapid evolution would strengthen inhibitory priority effects, making local species coexistence difficult. In contrast, Lankau (2009) and Vasseur et al. (2011) suggested that rapid evolution along a trade-off between intra-and inter-specific competitive ability would facilitate local species coexistence. Here we ask whether a similar mechanism can maintain regional diversity in a metacommunity with local inhibitory priority effects.

## Empirical motivation

In this study, we focus on inhibitory priority effects via interspecific interference, of which there are many empirical examples in microbes, animals, and plants. Microbes inhabiting floral nectar, for example, appear to change the chemical properties of nectar in a way that makes it harder for other, late-arriving species to colonize (Peay et al. 2012; Vannette et al. 2013). This type of self-serving habitat modification causes inhibitory priority effects. Similarly, in marine soft-bottom sediments, ghost shrimps and bivalves each modify grain size and oxygen content, and each group thrives better in its self-modified environment (Peterson 1984; Knowlton 2004), another case of inhibitory priority effects via interference. In plant communities, local positive feedbacks have been found to operate in some landscapes with interspersed patches of forest and heathland, mediated in this case by fire frequency and nutrient cycling (Petraitis and Latham 1999; Odion et al. 2010). More generally, many species of microbes and plants engage in “chemical warfare” with their competitors, causing inhibitory priority effects by interference.

In most of these cases, the producing organisms have resistance to their own chemicals. Some bacteria, for example, produce bacteriocins, compounds that inhibit or kill closely related strains or species, but do not affect the producing strain itself (Riley 1998). Many plants, including invasive species, produce allelopathic chemicals that harm heterospecific individuals more than conspecifics (Bais et al. 2003; Callaway and Ridenour 2004). Priority effects can also be caused by direct interference between heterospecific individuals. For example, some species of bacteria use contact-dependent growth inhibition (Ruhe et al. 2013), e.g., the so-called type VI secretion system, to inject toxic proteins directly into the cells of neighboring individuals, with bacteria generally resistant to the toxins produced by their own strain (Borenstein et al. 2015).

Empirical evidence also suggests that traits involved in inhibitory priority effects often evolve rapidly along a trade-off with other aspects of fitness. For example, rapidly evolving microbial resistance to bacteriocins or antibiotics often comes at a cost such as reduced growth rate (Riley 1998), reduced competitive ability (Gagneux et al. 2006), or “collateral sensitivity” to other types of antimicrobials (Pal et al. 2015). Similarly, in some plants, such as species of *Brassica,* both allelotoxin production and growth rate can evolve rapidly, but along a trade-off between the two traits (Lankau 2008; Lankau et al. 2009; Lankau 2011).

## Model overview and basic assumptions

Inspired by these empirical examples, we build a simple two-species metacommunity model with interspecific interference, which may arise, for example, via production of toxins that are harmful to members of the other species, but not to conspecifics. We consider a landscape that contains so many patches that stochasticity at the regional level can be neglected. Each patch has space for *k* individuals (*k* ≥ 2) and is always fully occupied. Generations are discrete and non-overlapping.

Interference occurs only among individuals living in the same patch in the same generation. Therefore, an individual’s fitness depends only on the current composition of the local patch community. There are no legacy effects, for example of toxins produced by previous generations. This is realistic for direct interference and also for many types of habitat modification, for example for toxins that rapidly decay or diffuse away.

Based on the empirical examples discussed above, changes in the composition of the metacommunity might lead to evolutionary change in the strength of interference on other species or in the resistance to interference from other species. In this study, we focus on the second possibility. We assume that all individuals have the same strength of interference, e.g., the same rate of toxin production, but differ in their sensitivity to heterospecific interference. Specifically, in each species, there are two types, one that is sensitive to interference by the other species and one that is completely or partially resistant but pays a cost *c* for this resistance. In a patch where the other species has frequency *q*, sensitive individuals of the focal species have relative fitness 1 — *d_s_ • *q**, where *d_s_* is a damage parameter for sensitive individuals, and resistant individuals have fitness 1 − *c − d_r_·q,* where *d_r_* < *d_s_* is the damage parameter for partially resistant individuals. With *d_r_* = 0, we have full resistance. Resistance evolves according to a haploid single-locus model with a mutation probability u per individual per generation. That is, with probability *u* an offspring of a resistant individual is sensitive and vice versa.

Assuming resistance is costly, sensitive individuals are favored if the other species is absent in the patch or at low frequency. In addition, we assume that (partially) resistant individuals are favored when the other species has a high local frequency. This is the case if

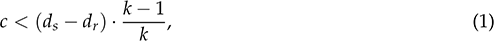

i.e., if the costs of resistance are small enough to make it worthwhile to invest in resistance when surrounded by heterospecific individuals.

Our goal is to explore whether ecologically similar species engaged in local interference can coexist due to rapid evolution alone, in the absence of other coexistence mechanisms. We therefore assume that parameters are identical across patches. Thus, there is no spatial environmental heterogeneity relevant to the coexistence of the species.

We will first consider a model in which there is global dispersal in every generation (no dispersal limitation) and then a model with dispersal limitation. The first model serves to explore the coexistence mechanism in its simplest form. The second model serves to demonstrate that this coexistence mechanism still operates under dispersal limitation and that metacommunities at an eco-evolutionary equilibrium can exhibit priority effects.

## Model with global dispersal

In this model version, all offspring produced in one generation are combined in a regional disperser pool. This regional pool is internal rather than external (*sensu* Fukami 2005) because its composition depends entirely on the cumulative local dynamics in the metacommunity. At the beginning of the next generation, the patches are recolonized according to the frequencies of the four types (two species, each with a sensitive and a resistant type) in the regional pool. Specifically, we assume that every spot in a patch is independently assigned to one of the four types such that local patch compositions follow a multinomial distribution. After recolonization, the individuals within a patch interact and then produce offspring according to the fitness values given above. Finally, the combined offspring from all patches make up the new regional disperser pool, thereby closing the life cycle.

Since we assume that the number of patches is very large, the metacommunity dynamics are fully specified by a deterministic model linking the frequencies of the four types in the regional disperser pool in successive generations. Let *p*1,*_r,t_* and *p*_1,*s,t*_ be the regional frequencies of resistant and sensitive individuals of species 1 at time *t*, and analogously for *p*_2,*r,t*_ and *p*_2,*s,t*_. We have *p*_1,*r,t*_ + *p*_1,*s,t*_ + *p*_2,*r,t*_ + *p*_2,*s,t*_ = 1. We assume that all patches contribute equally to the regional pool, for example because there is a fixed amount of resources in a patch. Thus, an individual’s contribution to the regional pool is its fitness divided by the summed fitnesses of all individuals in the patch. Such a selection regime is called “soft selection”. An alternative “hard selection” scenario where individuals contribute to the regional pool directly in proportion to their fitness values, i.e. independently of their patch neighbors, is explored in Online Appendix A.1.

### Invasion criteria

If each species, when rare, can invade a landscape dominated by the other species, we can expect stable species coexistence. We now check whether and under what conditions this “mutual invasibility” condition is fulfilled in our interference model. First, note that when one species is absent in the landscape, the sensitive type will be favored in the “resident” species. Without mutations (*u* = 0), the resistant type would go extinct; with mutations, it will be maintained at a small equilibrium frequency 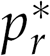 (mutation-selection balance).

For the rare species to increase in frequency, its members must have on average a higher fitness than their patch co-inhabitants. We first assume that mutation rate is negligible. All individuals of the resident species are then sensitive. Therefore members of the incoming rare species always share their patch with *k* – 1 sensitive resident individuals who are now exposed to interference by one heterospecific individual and therefore have fitness 1 – *d_s_* /*k*. Sensitive individuals of the incoming species have fitness 1 – *d_s_* (*k* – 1)/*k*, which is always smaller. Therefore the sensitive type of the rare species cannot increase in frequency. Resistant individuals of the incoming species have fitness 1 – *c* – *d_r_* (*k* – 1)/*k,* which is larger than the fitness of the resident individuals if

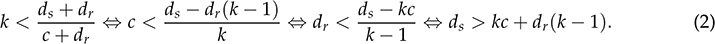

Thus for appropriate parameter combinations there is mutual invasibility and the two species will coexist regionally even if they interfere with each other locally. The conditions in (2) suggest that this “eco-evolutionary buffering” is facilitated by small local patch sizes, a cheap and efficient resistance mechanism, and a high interference damage in sensitive individuals. Note that the condition for the cost of resistance, c, is stronger than the trade-off assumption (1). The exact invasion criteria with *u* > 0 can be computed numerically (see Online Appendix A.2). For small *u*, (2) gives good approximations (Fig. A1).

Mutual invasibility requires genetic variation within species, i.e. the existence of both sensitive and resistant types. To see this, consider a modified model with only one type per species. We can even allow the species to differ in their trait values such that in a patch where species 1 has frequency *p* and species 2 frequency *q*, members of species 1 have fitness 1 – *c*_1_ – *d*_1_*q* and members of species 2 have fitness 1 – *c*_2_ – *d*_2_ *p*. For mutual invasibility, we need

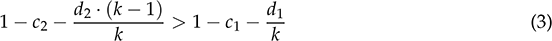

and

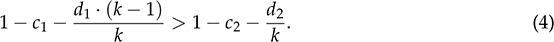

Summing inequalities (3) and (4) and simplifying, we obtain the condition *k* < 2, which is violating our additional assumption that *k* ≥ 2. Hence, mutual invasibility between monomorphic species in this model is not possible.

### Dynamics

While the above analysis tells us the conditions under which a regionally rare species can invade the landscape, it does not tell us how species will coexist and whether there are stable or unstable internal equilibria. To find out, we need to derive equations for the change in type frequencies over time. These equations will also allow us to explore whether a species that cannot invade when rare might be able to survive when it starts at higher initial frequency.

To derive the model equations under soft selection, we need to account for the contributions that patches of various composition make to the regional pool. We say a patch has configuration (*i*, *j*, *m*, *n*) if there are *i* species-1 resistant individuals, *j* species-1 sensitive individuals, *m* species-2 resistant individuals, and *n* species-2 sensitive individuals. Let *f_t_* (*i*, *j*, *m*, *n*) be the proportion of local patches with configuration (*i*, *j*, *m*, *n*) in generation *t*. Under the multinomial distribution,

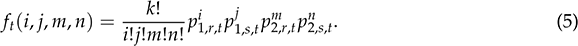

Offspring sent out by patches of the type (*i, j, m, n*) contains the four types in the following proportions:

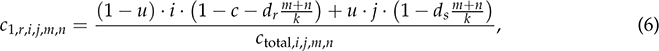

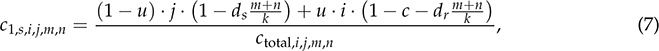

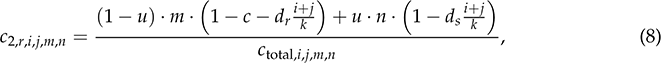

and

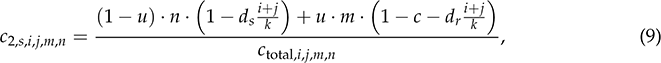

where *c*_total,*i,j,m,n*_ is the sum of the numerators of (6)–(9), ensuring that the contributions of the four genotypes sum to 1.

The new frequencies in the regional pool are then

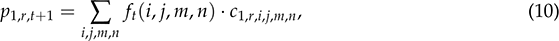

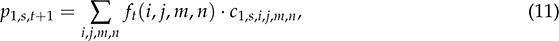

and analogously for species 2. Note that Σ*_i,j,m,n_* denotes a sum over all possible patch configurations, i.e. all possible combinations of *i*, *j*, *m*, *n* such that *i* + *j* + *m* + *n* = *k*. We now numerically iterate these equations to study the model with global dispersal in more detail.

### Critical frequencies

To derive the invasion criteria in (2), we assumed that the new species is so rare that its members always have a local abundance of 1. With a higher initial frequency, members of the new species may sometimes find themselves in a patch with one or more of their conspecifics and fewer heterospecific individuals and thus suffer less from interference. Therefore, we conjecture that a rare species may be able to invade if it starts above a certain critical frequency, even if it cannot invade from very low frequency. This would be an example for an Allee effect, specifically a strong demographic Allee effect, where the average per-capita growth rate is negative at low population density and increases with increasing density (Taylor and Hastings 2005).

To determine the critical frequency for a given parameter combination, we first let the frequencies of the two types in the resident population settle into mutation-selection balance. We then introduce the resistant type of the new species at larger and larger initial frequencies. The critical frequency is the smallest of our testing frequencies for which the new species increases in frequency over one generation.

As expected, the critical frequency is zero for all parameters fulfilling the mutual invasibility condition (Fig. 1). For each parameter, the critical frequency increases as we move further into the parameter range where the mutual invasibility condition is violated. For example, for local patch sizes, *k*, above the value in (2), the critical frequency increases with increasing *k* (Fig. 1 E).

**Figure 1:**
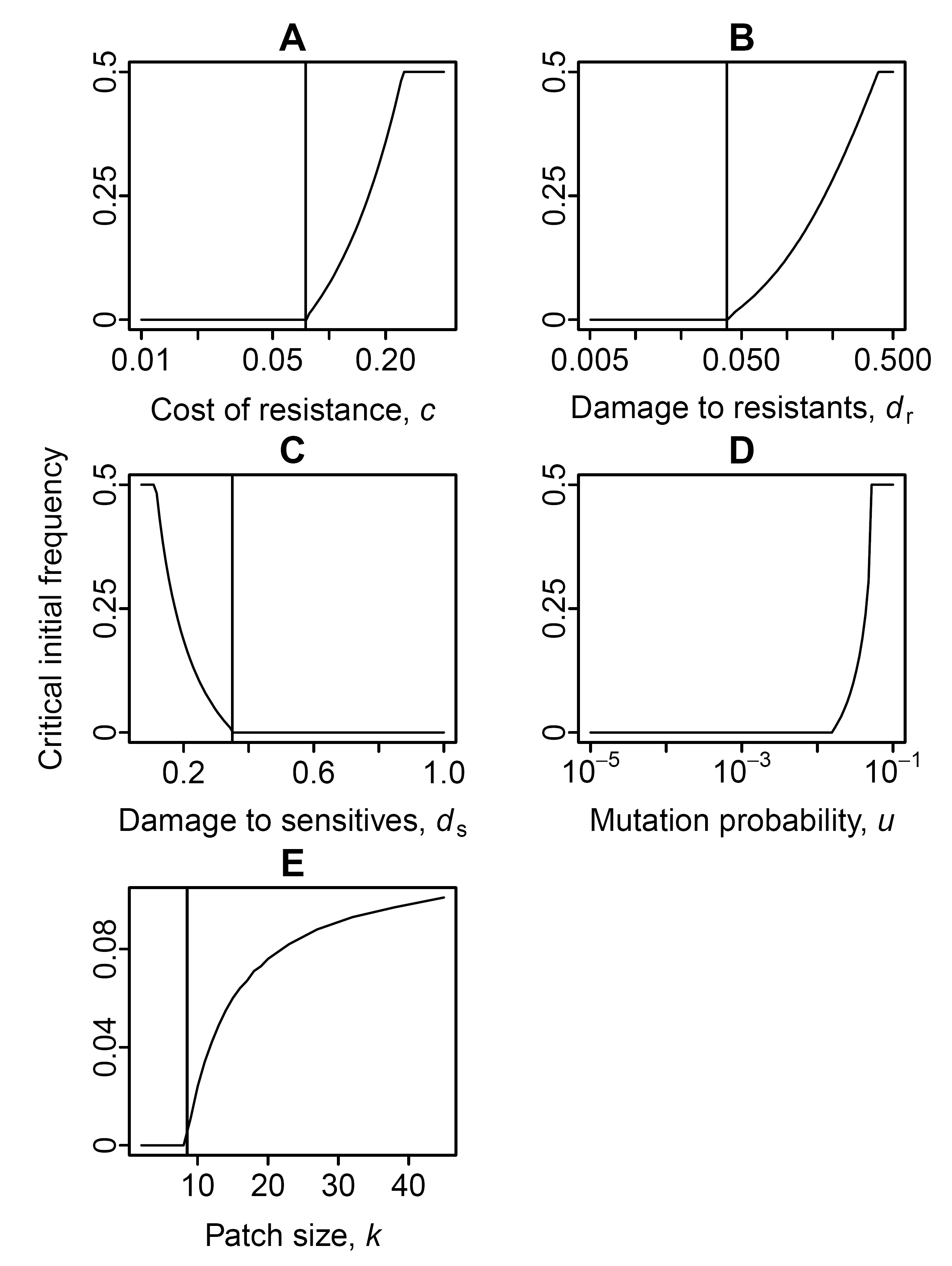
Critical frequency for the resistant type of a rare species to invade a resident population at mutation-selection equilibrium. If the critical frequency is zero, even an extremely rare species can invade and stable coexistence should be possible. The vertical lines indicate the approximate critical parameter values for the invasion of an extremely rare species (2). The underlying analytical argument did not take into account mutations and therefore the values may differ slightly from the numerically determined ones (see Fig. A1 for a comparison). All parameter combination fulfill the assumption (1). A critical frequency of 0.5 indicates that it was not possible for a rare species to invade. Default parameters: *k* = 6, *c* = 0.05, *d_s_* = 0.5, *d_r_* = 0.01, *u* = 0.001.

### Long-term behavior

Metacommunities in our model can exhibit five long-term behaviors: extinction of one species, symmetric coexistence at constant frequencies (Fig. 2 A), asymmetric coexistence at constant frequencies (Fig. 2 B), symmetric coexistence with fluctuating frequencies (Fig. 2 C), or asymmetric coexistence with fluctuating frequencies (Fig. 2 D). As discussed above, for some parameter combinations there is a critical frequency for invasion and depending on the initial conditions the long-term outcome will be either extinction of one species or one of the coexistence outcomes. Also, for each case of asymmetric coexistence, the initial conditions determine which of the two species will be the regionally common species in the long run.

**Figure 2:**
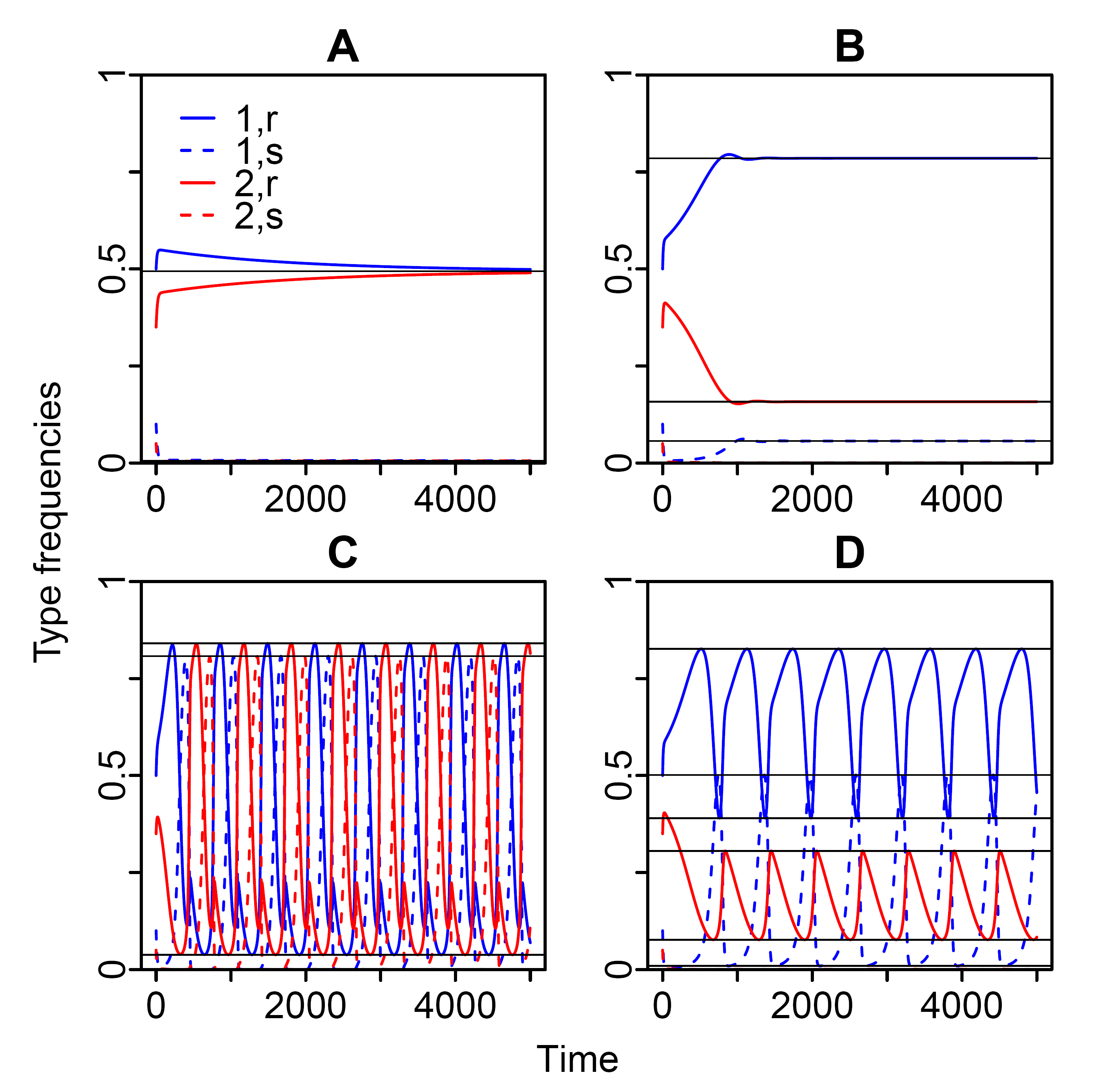
Time series for four parameter combinations illustrating the four types of coexistence outcome. (A) Symmetric coexistence at constant frequencies (*k* = 3, *d_r_* = 0.01). (B) Asymmetric coexistence at constant frequencies (*k* = 6, *d_r_* = 0.01). (C) Symmetric coexistence with fluctuations (*k* = 6,*d_r_* = 0.035). (D) Asymmetric coexistence with fluctuations (*k* = 12,*d_r_* = 0.01). Other parameters: *c* = 0.05, *d_s_* = 0.5, *u* = 0.001. Horizontal lines indicate minima and maxima along the cycle for the four types.

To systematically explore the role of the model parameters, we summarize the long-term behavior in terms of the minimum and maximum frequency for each of the four types along the cycle (see horizontal lines in Fig. 2). Without fluctuations, minimum and maximum are the same. Fig. 3 explores the influence of the five model parameters on the long-term behavior. With changing parameter values, the system typically goes through a series of qualitatively different outcomes. For example with increasing damage parameter for resistants (Fig. 3 B), we first observe symmetric coexistence without fluctuations, then asymmetric coexistence without fluctuations, then asymmetric coexistence with increasingly large fluctuations, then symmetric coexistence with fluctuations, and finally coexistence is no longer possible.

**Figure 3:**
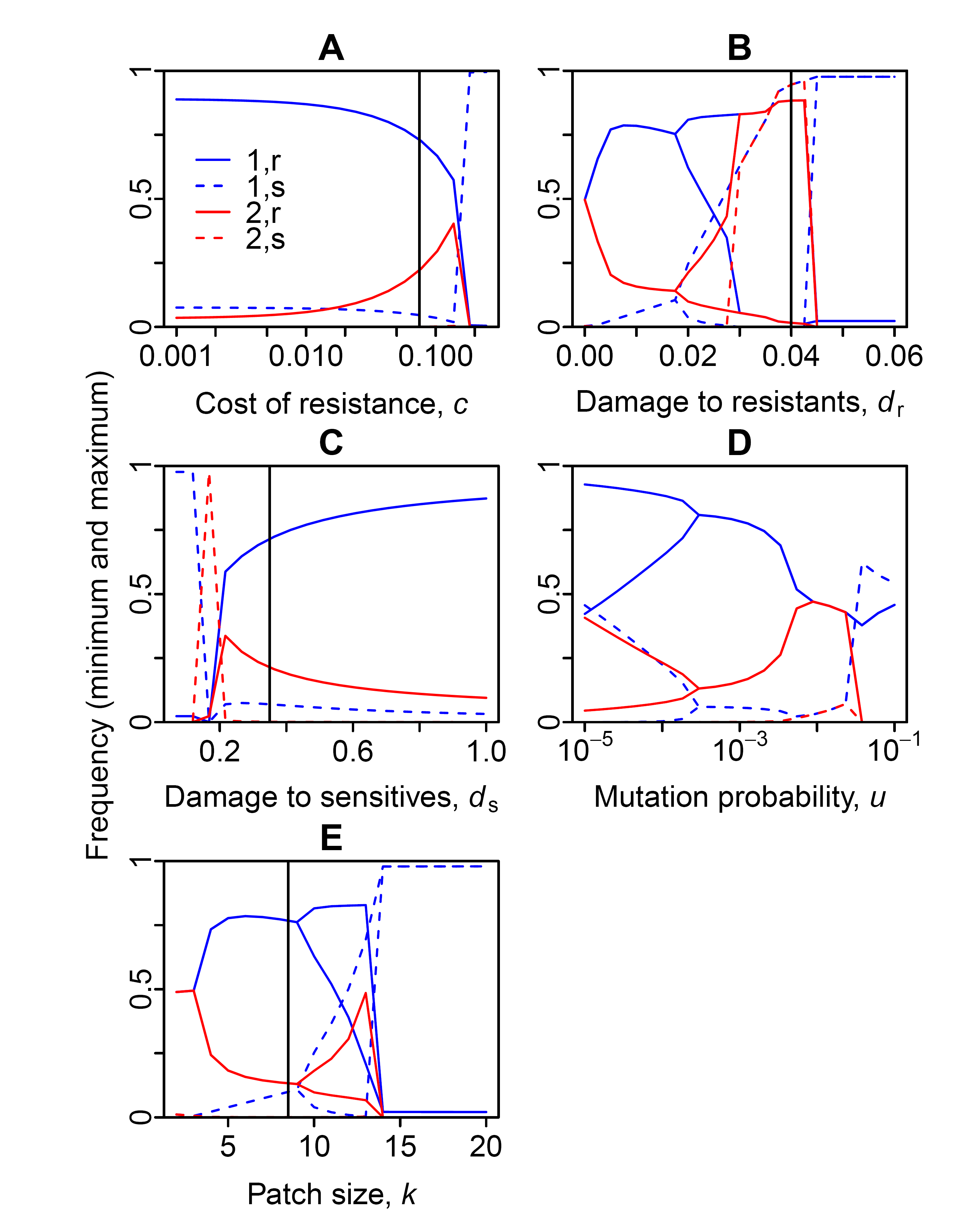
Minimum and maximum type frequencies along the respective attractor as a function of (A) the cost of resistance, *c*, (B) the maximum interference damage in partially resistant individuals, *d_r_*, (C) the maximum interference damage in sensitive individuals, *d_s_*, (D) the mutation probability, *u*, and (E) the local patch size, *k*. In each panel the respective other four parameters take the following default values: *k* = 6, *c* = 0.05, *d_s_* = 0.5, *d_r_* = 0.01, *u* = 0.001. Initial conditions: *P*_1,*r,0*_ = 0.6, *p*_2,*r*,0_ = 0.4, *p*_1,*s*,0_ = *p*_2,*s*,0_ = 0. Vertical lines indicate the critical parameter value for mutual invasibility.

Overall the role of the parameters is consistent with the above results on mutual invasibility. Increases in *k*, *c*, or *d_r_* destabilize coexistence, whereas an increase in *d_s_* facilitates coexistence. However, both with decreasing *c* and increasing *d_s_*, coexistence becomes more and more asymmetric. Although such coexistence may always be stable in the deterministic system, one of the species may go extinct rapidly in metacommunities of finite size with regional stochasticity. In such stochastic metacommunities, intermediate values of *c* and *d_s_* might be most conducive to long-term coexistence. Also for the mutation probability, *u*, intermediate values appear most conducive to coexistence. On the one hand, too much mutational noise prevents species from adapting to the current metacommunity state. With too few mutations, on the other hand, fluctuations in regional frequencies are large, such that the rare species would have a large extinction risk in a metacommunity of finite size.

## Model with dispersal limitation

We now introduce some more permanent spatial structure. So far, patches were fully recolonized in each generation. Now this only happens with probability *e* per patch and generation. Otherwise, each member of the local patch in the new generation is drawn either from the regional pool with probability μ, the dispersal probability, or is the offspring of a local individual with probability 1 – *μ*. All species have the same dispersal ability. The smaller *μ* and *ϵ* are, the more permanent is the spatial structure in the landscape. By setting *ϵ* = 1 or *μ* = 1 or both, we recover the model with global dispersal as a special case.

With dispersal limitation, it is no longer sufficient to trace the regional frequencies of the four types. We need to keep track of the proportion of patches *f_(i,j,m,n),t_* in the metacommunity for each of the possible patch configurations (*i*, *j*, *m*, *n*) with *i* + *j* + *m* + *n* = *k*. Over a single generation without full recolonization, a patch with configuration (*i*′, *j*′, *m*′, *n′*) at time *t* turns into a patch with configuration (*i*, *j*, *m*, *n*) at time *t* + 1 with probability

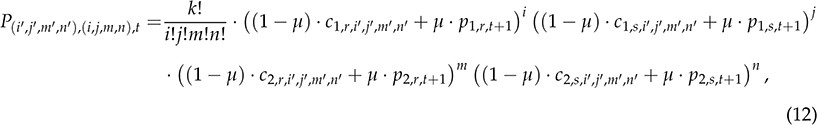

which is a multinomial distribution. The frequencies in the regional pool, *p*_1,*r,t*+1_, *p*_1,*s,t*+1_, *p*_2,*r,t*+1_, and *p*_2,*s,t*+1_ are given by (10) and the analogous equations for the other types. Finally,

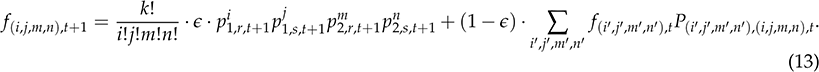

### Coexistence with dispersal limitation

Numerical iterations indicate that dispersal limitation enhances eco-evolutionary buffering and thus stabilizes coexistence (Fig. 4). Compared to the case with global dispersal (see Fig. 3), dispersal limitation leads to coexistence, and especially symmetric coexistence, over a wider range of values for the parameters *c*, *d_r_*, *d_s_*, and *u* (Fig. 4 A-D). We did not attempt to increase the local community size, *k*, under dispersal limitation because the number of patch types to be traced rapidly increases with *k*, which makes computations unfeasible. Even for parameter combinations that do not allow for coexistence with global dispersal (dispersal probability *μ* = 1 or recolonization probability *ϵ* = 1), a decrease in dispersal and recolonization probabilities can make coexistence possible (Fig. 4 E-F). With decreasing dispersal and recolonization probabilities, we first observe fluctuating coexistence and eventually also symmetric coexistence without fluctuations.

**Figure 4:**
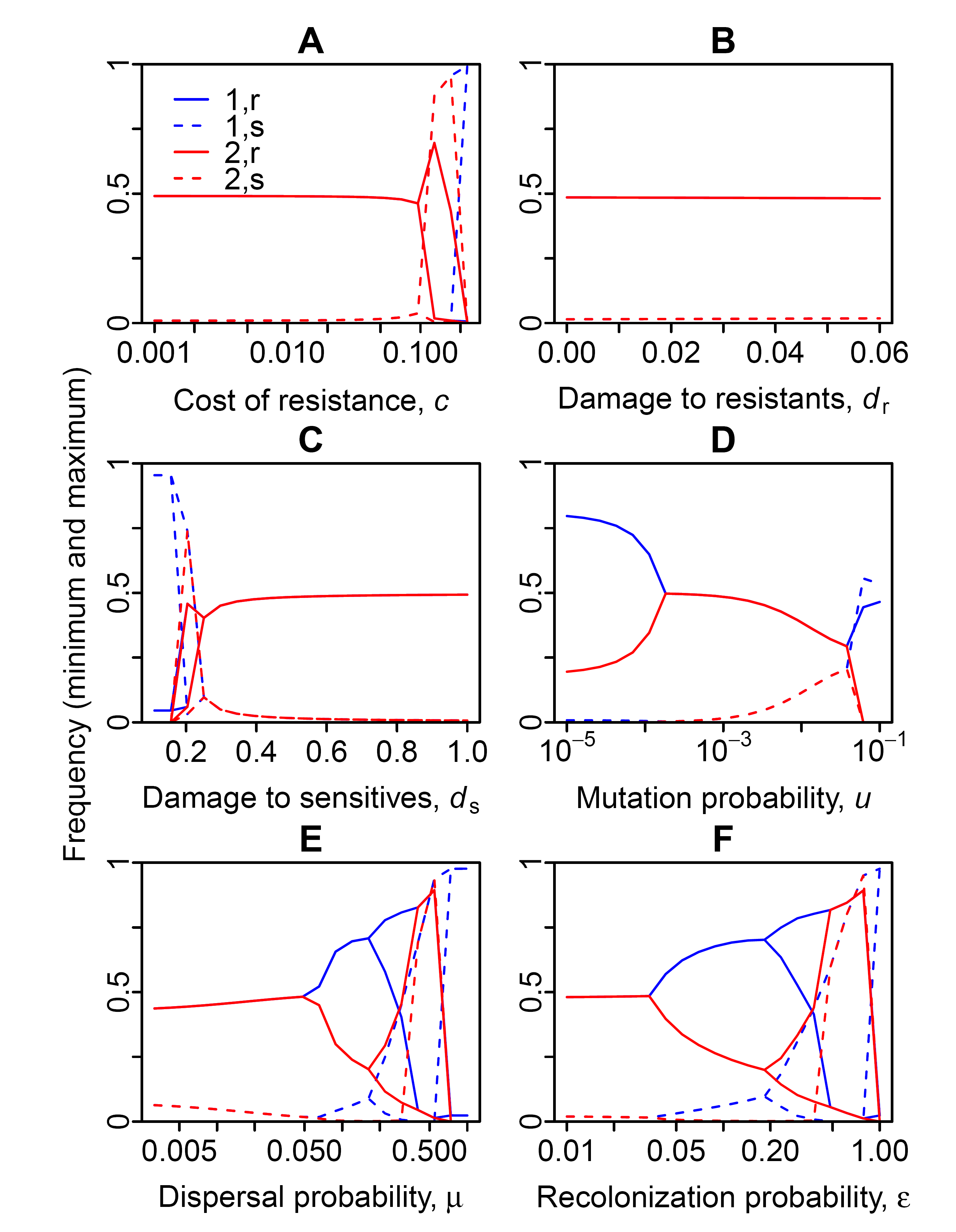
Minimum and maximum type frequencies along the respective attractor with dispersal limitation as a function of (A) the cost of resistance, *c*, (B) the maximum interference damage in partially resistant individuals, *d_r_*, (C) the maximum interference damage in sensitive individuals, *d_s_*, (D) the mutation probability, *u*, (E) the dispersal probability, *μ*, and (F) the recolonization probability, e. In (A–D) *d_r_* = 0.01 and in (E–F) *d_r_* = 0.05 to show a broader range of behaviors. In each panel the respective other parameters take the following default values: *k* = 6, *c* = 0.05, *d_s_* = 0.5, *u* = 0.001, *m* = 0.05, *e* = 0.02. Initial conditions: p_1,*r*,0_ = 0.6, p_2,*r*,0_ = 0.4, p_1,*s*,0_ = p_2,*s*,0_ = 0.

### Priority effects

We have shown so far that eco-evolutionary buffering can lead to the regional coexistence of species engaged in local interference competition and that this works even better with dispersal limitation. But the central question for the purpose of this study is whether or not we still observe priority effects in these metacommunities. To address this question, we define priority effects as cases with positive local frequency dependence. That is, we have a priority effect if a locally rare species tends to decrease in local frequency whereas a locally common species tends to increase in frequency. The motivation for this definition is that positive frequency dependence helps a species that is common among the initial colonizers to defend the patch against later invasions by the respective other species.

To determine whether or not there are priority effects at some time point *t*, we thus need to characterize the local dynamics. For this, we first consider all possible patch configurations (*i*, *j*, *m*, *n*) and compute the expected population size of species 1 after one generation

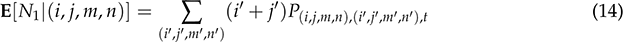

without full recolonization. Given the current state of the metacommunity as specified by the proportions of patches with the various configurations, *f_(i,j,m,n),t_*, we then compute E[Δ_*l*_], the expected change in the local population size of species 1 in patches with current population size *l*:

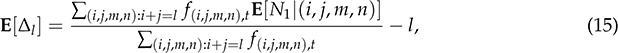

where the fraction represents a weighted average of the expectations in (14) over all patch types that have *l* individuals of species 1 (taking together the sensitive and the partially resistant type) with the weight proportional to the frequency of the respective patch types.

Under neutrality (*d_s_* = *d_r_* = *c* = 0),

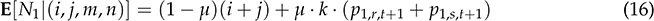

and thus

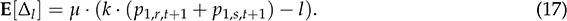

The expected change is positive whenever the local frequency is below the regional frequency, and negative when the local frequency is above the regional frequency. Local communities thus tend to become more similar in composition to the regional pool because the incoming migrants reflect the regional frequencies. Without dispersal (*μ* = 0), the expected change in local species abundances would be zero under neutrality (E[Δ_*l*_] = 0 for all *l*).

We now formalize the notion of positive frequency dependence, and say that there is a priority effect if E[Δ_*l*_] < 0 for all 0 < *l* < *k*/2 and E[Δ_*l*_] > 0 for all *k*/2 < *l* < *k*. The states 0 and *k* are not taken into account because an absent species cannot decrease any further. Neither do we take into account the expected change at local population size *k*/2 (which only exists for even patch sizes, *k*). Note that it is sufficient to check the priority-effect conditions for species 1 because the expected change in the local population size of species 2 is just – E[Δ_*l*_]. Thus if species 1 is expected to increase when common and decrease when rare this is necessarily true also for species 2. Note also that this is a rather strict definition of a priority effect because even if the condition is not fulfilled there is local interference and species that are initially common may generally defend patches for longer than under neutrality. However, to be conservative and highlight the clear-cut cases, we use the stricter definition.

In Fig. 5, we give examples for the expected local dynamics under different types of coexistence outcome and compare them to the expected local dynamics under neutrality, i.e. with *d_s_* = *d_r_* = *c* = 0. The top row shows an example of symmetric coexistence without fluctuations (Fig. 5 A), whose local dynamics fulfill our priority-effect criterion (Fig. 5 B). That is, the expected change in the local population size of species 1 is negative (i.e. species 2 is expected to increase) if there are 1 or 2 members of species 1 in a patch of size 6, and positive when there are 4 or 5 members. The condition for priority effects is fulfilled both at equilibrium (purple lines) and at an earlier time point where species 1 is more common in the landscape (green lines). At equilibrium, the situation is entirely symmetric, with an expected change of zero when both species have local abundance 3. At the earlier time point, the local dynamics are slightly asymmetric with the regionally common species having a slight local disadvantage. For example in patches with the same number of individuals of both species, species 1 is expected to slightly decrease in population size. This disadvantage of species 1 was due to a higher frequency of the sensitive type. This example illustrates how stable regional coexistence is possible even with local priority effects. When the system is perturbed and one species becomes more common, it will soon experience an increase in the frequency of sensitive individuals. This will make the local dynamics slightly asymmetric in favor of the regionally rare species, thus allowing it to recover.

**Figure 5:**
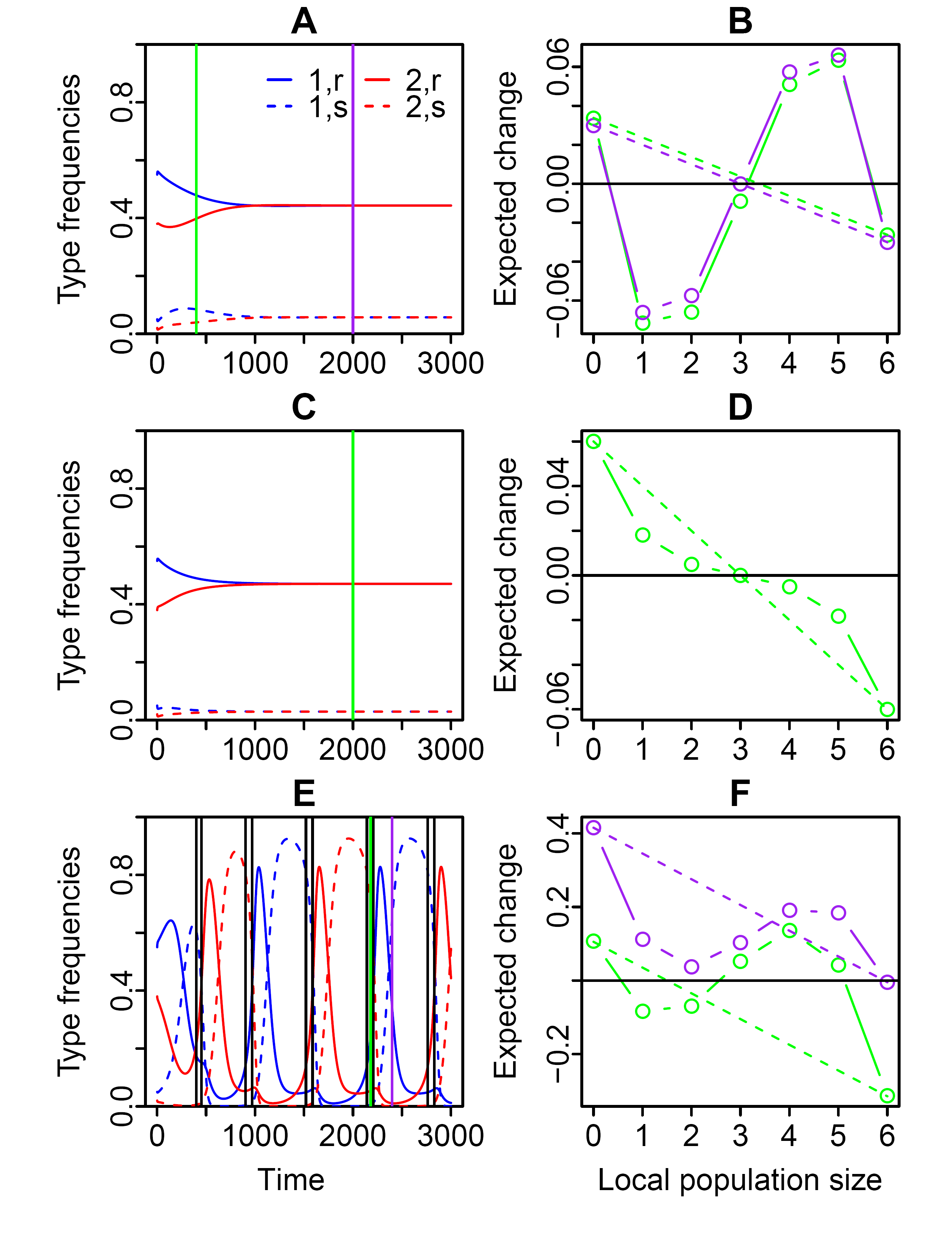
Examples for regional dynamics (left column) and local feedbacks (right column) under dispersal limitation. The panels in the right column show the expected change in the local population size of species 1 (symbols and solid lines) and compare it to the expected change under neutrality (*d_s_* = *d_r_* = *c* = 0, dashed lines). The different colors in the right column correspond to different times, indicated by vertical lines in the left column. Parameter values: (A, B) *d_r_* = 0.1, *μ* = 0.01, (C, D) *d_r_* = 0.01, *μ* = 0.02,(E, F) *d_r_* = 0.2, *μ* = 0.07. The black vertical bars in (E) indicate the start and end points of time periods with priority effects. Other parameters: *k* = 6, *d_s_* = 0.5, *c* = 0.05, *u* = 0.001, *ϵ* = 0.02.

The second row in Fig.5 has the same parameters as the first row except that the maximum damage to resistants, *d_r_*, is smaller, and dispersal probability, *μ*, is higher. Now local interference between species is not strong enough to counteract dispersal and local communities on average become more similar in composition to the regional pool. Thus, according to our definition, there are no priority effects. But note that the approach of local frequencies to regional frequencies is slower than under neutrality.

For coexistence outcomes with fluctuating regional frequencies, the transition probabilities between patch types (12) and therefore also the expected local dynamics change over time. In the example in Fig. 5 E and F, most of the time either species 1 or species 2 is the dominant competitor (e.g. at the purple time point). However, there are also brief time periods (between adjacent vertical lines in Fig. 5 E) where interference effects are relatively symmetric and the priority-effect condition is fulfilled.

More generally, whether or not there are priority effects in the long run depends on the strength of interference in relation to the dispersal probability, *μ* (Fig. 6). The total interference effects are the sum of interference effects on the sensitive types and interference effects on the partially resistant types. Under constant symmetric coexistence, the sensitive types of both species are rare, but if dispersal probability is small, interference effects on them can be sufficient to cause priority effects even if the other types are completely resistant (*d_r_* = 0). For larger dispersal probabilities, priority effects only occur when the partially resistant types are still sufficiently sensitive, i.e. if *d_r_* is sufficiently large. In the region of parameter space with symmetric constant coexistence, the local dynamics stay constant over time, and therefore priority effects are either present all the time or absent all the time. There is a small region of parameter space with asymmetric or symmetric fluctuating coexistence where priority effects are present part of the time. However, most parameter combinations with asymmetric coexistence or symmetric fluctuating coexistence do not exhibit priority effects.

**Figure 6:**
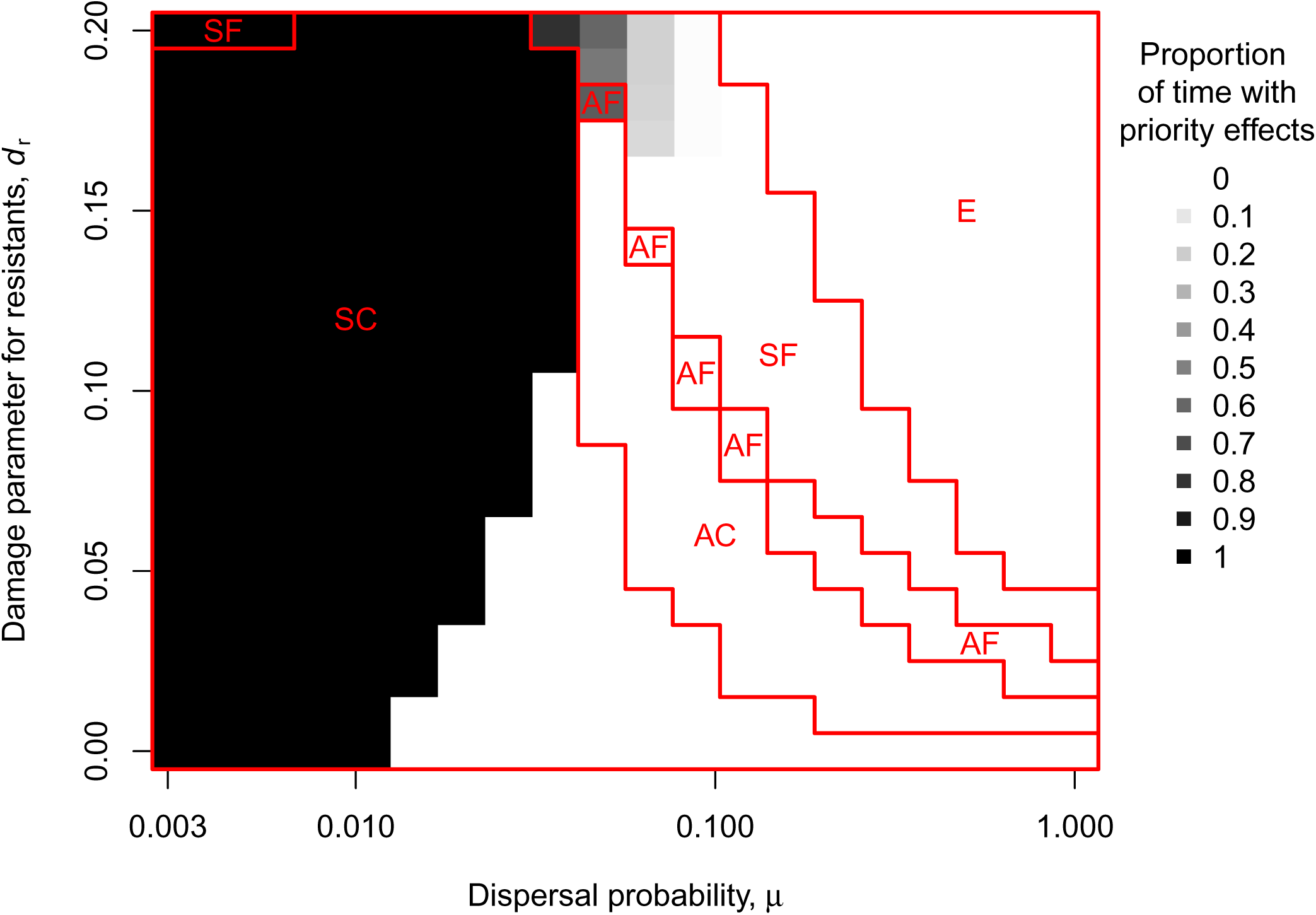
Proportion of times during which there are clear priority effects as a function of the dispersal probability, *μ*, and the damage parameter of partially resistant individuals, *d_r_*. The red lines and labels indicate regions in parameter space with the various outcome types (SC: symmetric constant coexistence, AC: asymmetric constant coexistence, AF: asymmetric fluctuating coexistence, SF: symmetric fluctuating coexistence, E: extinction of one species). Other parameters: *k* = 6, *d_s_* = 0.5, *c* = 0.05, *u* = 0.001, *ϵ* = 0.02. Initial conditions: *p*_1,*r*,0_ = 0.6, *p*_2,*r*,0_ = 0.4, *p*_1,*s*,0_ = *p*_2,*s*,0_ = 0.

## Discussion

Taken together, our results suggest a new “eco-evolutionary buffering” hypothesis for species coexistence in the presence of local priority effects. In this hypothesis, we assume that resistance to heterospecific interference is costly, such that the strength and direction of selection on resistance depend on regional relative frequencies of species. Thus, when one species becomes regionally rare, resistance against this species does not pay off any longer for members of the other species. The resulting loss of resistance can then be exploited by the rare species to recover. Consequently, both regional species diversity and intra-specific genetic variation will be maintained, even though local priority effects may persist. Our focus on priority effects and coexistence at the regional rather than local scale is the main novelty of our study compared to previous work on similar coexistence mechansisms (Levin 1971; León 1974; Pease 1984; Lankau 2009; Vasseur et al. 2011). Under eco-evolutionary buffering, the parameter combinations that allow for the most stable coexistence (symmetric without fluctuations) are also those most likely to maintain priority effects. In these cases, coexistence only requires that priority effects become slightly asymmetric, with the regionally common species less likely to take over a patch from the regionally rare species than vice versa.

### Requirements for eco-evolutionary buffering

In addition to interspecific interference, there are several requirements for eco-evolutionary buffering. One is intra-specific genetic variation in resistance to interference by other species. Without such variation, species do not coexist in our model. We have assumed that variation in resistance is due to two alleles at a single locus. However, similar coexistence mechanisms for single communities can work with quantitative traits (Pease 1984; Vasseur et al. 2011).

A second requirement is a trade-off between resistance to interference and maximum fitness. We have found that eco-evolutionary buffering works particularly well if the resistance mechanism is efficient (small *d_r_*) and not very costly (small *c*). In a broad sense, the trade-off underlying eco-evolutionary buffering can be regarded as a competition-colonization trade-off, where the roles are assigned depending on which species is more common in the region. The regionally common species corresponds to the better colonizer because it is more likely to have the majority in newly colonized patches and thus benefits more often from priority effects. The regionally rare species is better at invading patches dominated by the other species and thus corresponds to the better competitor.

A third requirement is stochastic variation in local community composition, for which small local community size, *k*, is required. To understand this requirement, it helps to consider a metacommunity with one species that is regionally very rare and the other very common. If local community size is small, most members of the common species will be in patches without a single member of the rare species and are hence selected to lose resistance. Members of the rare species, on the other hand, have a local frequency of at least 1/*k*. Since interference damage is proportional to local frequency, the rare species can do more damage with smaller *k*. Other studies on competitive metacommunities have also found that local community size affect coexistence, but sometimes with opposing results. For example, Orrock and Watling (2010) studied the regional coexistence of two species under a competition-colonization trade-off and showed that large local community size facilitated regional coexistence because they made the local dynamics and thus patch-take overs by the better competitor more deterministic. However, they assumed that the initial frequency of a patch invader was independent of local community size, which we did not assume in this study.

A fourth requirement is a large number of local patches. There will otherwise be considerable stochasticity in regional abundance, making the extinction of genotypes and species likely, even in cases where the deterministic model predicts stable coexistence. Also, regional stochasticity may cause species to fall below the critical frequency in cases where coexistence is possible but the mutual invasibility condition is not fulfilled.

A fifth requirement is that competitive interactions are local, which is fulfilled if there is dispersal limitation. Without dispersal limitation, it is fulfilled as long as all patches contribute equally to the regional pool, such that individuals contribute more if the other individuals they share the patch with are less fit. In Online Appendix A.1, we consider an alternative scenario where individuals contribute in proportion to their fitness, independently of the other patch inhabitants. Under this assumption mutual invasibility is not possible. The two scenarios are referred to as soft selection vs. hard selection. The former is generally more conducive to the maintenance of diversity (Christiansen 1975).

Finally, extinction-recolonization events are required for local priority effects to be observed in metacommunities with eco-evolutionary buffering. Without such disturbance or when disturbance occurs at a smaller scale than local positive feedbacks, the landscape may settle into a configuration where each patch is dominated by one species. The regional dynamics then come to a halt and species can coexist for extended periods of time, as demonstrated by Molofsky et al. (1999, 2001) and Molofsky and Bever (2002) for spatially explicit models. Since there is no disturbance to initiate new rounds of local community assembly, priority effects will no longer be operating.

Although these requirements may seem stringent, they may be fulfilled in many real communities, particularly those of sessile animals in intertidal habitats, herbaceous plants in small patches such as tussock islands, and parasite or parasitoid insects that co-infect hosts (Levine 2000; Mouquet and Loreau 2002; Fukami and Nakajima 2013; Zee and Fukami 2015). For exam-ple, in metacommunities of co-infecting parasitic flatworms, local community sizes in a single host individual are often small, e.g. on the order of ten for fish eye flukes (Seppala et al. 2009). Furthermore, there is evidence for inhibitory local priority effects (Leung and Poulin 2011). Interspecific interference among parasitic flatworms might often involve the host immune system (Seppala et al. 2009; Leung and Poulin 2011), but in some cases, there are also specialized soldier individuals that kill new individuals attempting to invade the same host individual (Hechinger et al. 2011). Some of these interference effects are strain-specific and have been suggested to maintain genetic variation within parasite species (Seppala et al. 2009). Even in some microorganisms, relevant interaction neighborhoods may consist of few individuals, for example in highly structured bacterial biofilms (Cordero and Datta 2016). An important form of interference in such biofilms is contact-dependent growth inhibition where individuals attach to neighboring cells and inject toxins (Ruhe et al. 2013).

One main difference between eco-evolutionary buffering and previous models for coexistence in evolving metacommunities concerns environmental heterogeneity. Eco-evolutionary buffering requires that individuals experience different patch community compositions due to intrinsically generated and stochastic variation, but the environmental conditions can be the same in all patches. By contrast, previous models for coexistence in evolving metacommunities require extrinsically generated spatial environmental heterogeneity. Of particular relevance here is the work on evolutionary monopolization (De Meester et al. 2002; Urban 2006; Urban et al. 2008; Urban and De Meester 2009). If evolution is fast enough relative to migration, populations can locally adapt to the various patch environments and prevent later-arriving species from invading. Species can then also coexist in the landscape but only as long as the patches are not disturbed and recolonized. Another aspect of spatial structure that is not necessary for coexistence via eco-evolutionary buffering is dispersal limitation. However, dispersal limitation facilitates coexistence and is required for local priority effects to be observed. Thus unlike similar eco-evolutionary coexistence mechanisms which are sometimes destabilized with increasing spatial structure (Vellend and Litrico 2008; Lankau 2009), eco-evolutionary buffering becomes even more stable with more persistent spatial structure.

### Future directions

This study is only a first step toward understanding the role of eco-evolutionary buffering in the maintenance of species diversity. In future research, it would be useful, for example, to consider evolution of toxin production or other forms of interference in addition to evolution of resistance. Whereas a resistance mutation can directly reduce the death rate of the mutated individual, a toxin-production mutation first reduces the fitness of heterospecific individuals. Indirectly, it may then benefit the mutated individual, but also other conspecific individuals that do not pay the fitness cost. Hence interference can be an altruistic trait and its evolution can be affected by cheating. It remains unclear how readily eco-evolutionary buffering occurs in these circumstances. Other questions that should be addressed include: whether eco-evolutionary buffering works with more than two interacting species; whether it works for diploid sexual organisms; and how eco-evolutionary buffering interacts with spatial and temporal environmental heterogeneity to affect regional coexistence.

Besides the eco-evolutionary buffering mechanism we have studied here, a number of other mechanisms could potentially buffer regional diversity in the presence of priority effects. These mechanisms warrant further investigation. First, simple patch-occupancy models suggest that, by virtue of spatial structure alone, two identical competitors can coexist in a region even if there is some local inhibition (Slatkin 1974; Hanski 1983). However, this requires doubly-occupied patches to send out the same number of colonists of both species (Taneyhill 2000), an assumption that has been criticized as giving an “unfair” advantage to the regionally rare species (Wang et al. 2005). Second, a predator that forages at a regional scale may either exhibit behavioral plasticity or evolve rapidly to preferentially prey on regionally common species (e.g., Hughes and Croy 1993). Third, if patches differ in environmental conditions, regionally rare species may be better at evolutionary monopolization of patches (Urban and De Meester 2009; De Meester et al. 2016) as they suffer less from the inflow of maladapted migrants (Lankau 2011). Fourth, if individu-als experiencing strong interference can move to another patch, regional coexistence is possible (Ruokolainen and Hanski 2016) and it would be interesting to explore whether priority effects can also be maintained in this setting. Finally, at a long evolutionary time scale, any factor that accelerates speciation rate would help to maintain a speciose regional pool. Speciation rate itself may be affected by local priority effects (Fukami et al. 2007). Interactive effects of speciation and priority effects on the generation and maintenance of species pools are a particularly interesting topic for future research.

For empirical tests of eco-evolutionary buffering, one could choose two species engaging in interspecific interference, and for each species pick two genotypes such that one of them is more resistant to interspecific interference but the other one has a higher growth rate in the absence of the other species. As different treatments, one could initialize a patchy landscape either using only one genotype per species or using a mixture of both genotypes for each species. Our theory predicts that long-term coexistence is impossible in the treatment with only one genotype per species but might be possible in the treatment with two genotypes per species. Such an experiment could be performed, for example, in the field with herbaceous plants inhabiting a landscape of tussock micro-islands (as in Levine 2000) or in the laboratory with parasites co-infecting a host population.

## Acknowledgments

For discussion and comments on an earlier stage of the project, we thank members of the community ecology group at Stanford, particularly Luke Frishkoff, Po-Ju Ke, Devin Leopold, Erin Mordecai, and Rachel Vannette, as well as Angela Brandt, Joachim Hermisson, Kotaro Kagawa, Mike McLaren, Akira Mori, Pleuni Pennings, Dmitri Petrov, and Chris Klausmeier and two anonymous reviewers. MJW acknowledges fellowships from the Stanford Center for Computational Evolutionary and Human Genomics (CEHG) and from the Austrian Science Fund (FWF, M 1839-B29). This work was also supported by the NSF (DEB 1149600) and Stanford University’s Terman Fellowship.

## Online Appendix A: Supplementary methods and results

### Online Appendix A.1 Modified model with hard selection

Under hard selection, individuals contribute to the regional pool in proportion to their fitness values. Thus patches with a high average fitness make a larger total contribution to the regional pool than patches with a low average fitness.

To obtain the model equations for hard selection, we need the average numbers of offspring 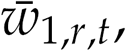 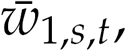 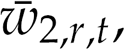 and 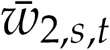 produced by individuals belonging to the four types. We have

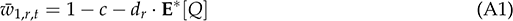

and

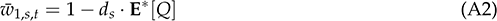

where 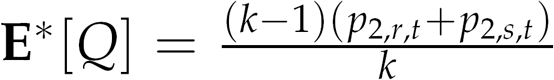 is the average local frequency of species 2 experienced by species 1 individuals. Note that this is lower than *p*_2,*r,t*_ + *p*_2,*s,t*_ because from the perspective of each species-1 individual, there are only *k* – 1 spots left that can potentially be occupied by species 2. This intuitive result can also be formally derived by averaging over all possible local patch configurations. The expressions for 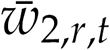 and 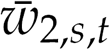 are analogous. The regional frequency of species-1 resistant individuals in the next generation is then

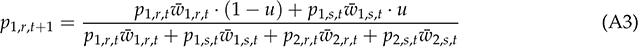

and analogously for those of the other types.

A rare species will be able to invade, i.e. increase in frequency, if its average fitness is larger than the average fitness of the resident population. The average fitness in the resident population at mutation-selection balance is 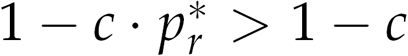 and when we now introduce the other species at very low frequency in the landscape, this value will not change by much. The members of the rare species will generally find themselves alone in a patch surrounded by *k* – 1 members of the other species. Hence the average fitness of these sensitive individuals is 1 – *d_s_*(*k* – 1)/*k*, which is smaller than 1 – *c* by assumption (1), and the average fitness of resistant individuals is 1 – *c – d_r_ (k –* 1)/k, which is also smaller than 1 – *c* (these values for the average fitnesses also follow from (A1) and (A2) by setting the frequency of the other species to 1). Therefore both types of the rare species have an average fitness below that of the resident population. The rare species cannot increase in frequency under hard selection.

### Online Appendix A.2 Invasibility conditions with a resident species at mutation selection balance

In the main text, the critical parameter values for mutual invasibility (2) were derived under the assumption that the resident population consists only of sensitive individuals. However, because of recurrent mutations, the resistant type will be present in the resident population at low frequency even if it is disfavored in the absence of the other species (mutation-selection balance). Here we explore how the estimates for the critical parameter values change if we account for these rare resistant individuals in the resident population.

To determine the equilibrium frequency of the resistant type in the resident species, say species 1, we set *p*_2,*r,t*_ = *p*_2,*s,t*_ = 0 and use (5), (6), (7), and (10) to obtain an equation for the equilibrium frequency, *p_r*_*, of the resistant type:

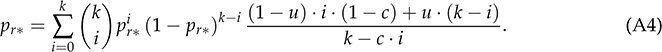

We then use the uniroot function in R to solve this equation numerically for *p_r*_*.

The condition for the other species to invade from very low frequency is that a single resistant individual of the other species contributes on average more than one *k*th of the offspring sent out by its patch, i.e.

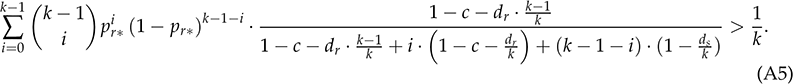

To obtain the value of a certain parameter at which the resident species becomes invasible, we again use the uniroot function to numerically solve this equation for the respective parameter of interest. In Fig. A1, the resulting critical values of the parameters *c*, *d_r_*, *d_s_*, and *k* are given as a function of the mutation probability, *u*. Unless the mutation rate is very large, the approximations in (2) in the main text are very accurate.

**Figure A1:**
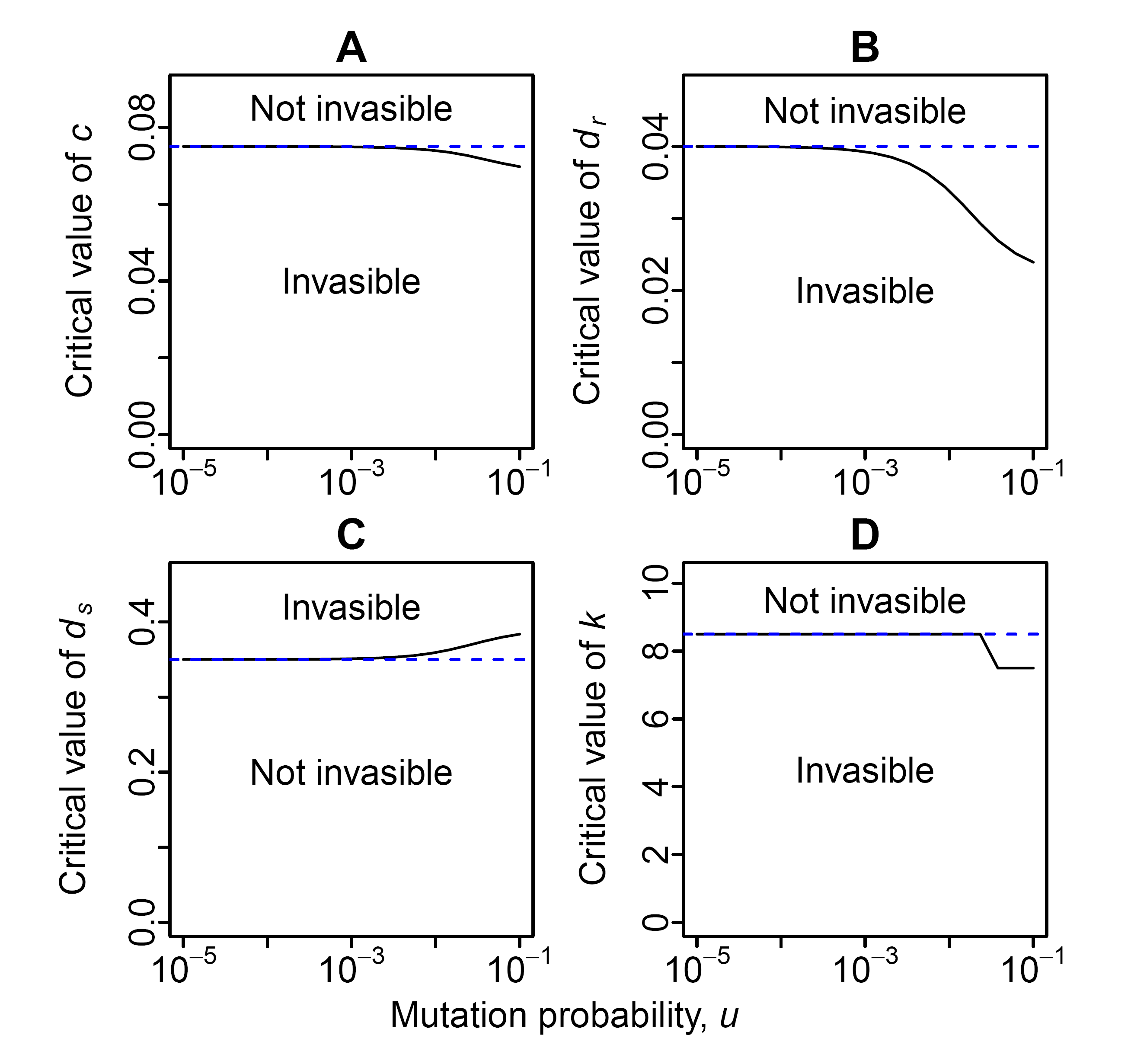
Critical parameter values for coexistence when taking into account the presence of the resistant type in the resident population at mutation-selection balance (black solid lines). Blue dashed lines indicate the approximations in (2), which assume that the resident population only consists of sensitive individuals. (A) Cost of resistance, (B) Damage to resistants, (C) Damage to sensitives, (D) Patch size. Default parameters: *k* = 6, *d_r_* = 0.05, *d_s_* = 0.5, *c* = 0.05.

